# Tracking selective rehearsal and active inhibition of memory traces in directed forgetting

**DOI:** 10.1101/864819

**Authors:** Marie-Christin Fellner, Gerd T. Waldhauser, Nikolai Axmacher

## Abstract

Selectively remembering or forgetting newly encountered information is essential for goal-directed behavior. It is still an open question, however, whether intentional forgetting is an active process based on the inhibition of unwanted memory traces or whether it occurs passively through reduced recruitment of selective rehearsal [1,2]. Here we show that intentional control of memory encoding relies on both, enhanced active inhibition and decreased selective rehearsal, and that these two processes can be separated in time and space. We applied representational similarity analysis (RSA [3]) and timefrequency analysis to EEG data during an item-method directed forgetting experiment [4]. We identified neural signatures of both the intentional suppression and the voluntary upregulation of item-specific representations. Successful active forgetting was associated with a downregulation of item-specific representations in an early time window, 500ms after the instruction. This process was initiated by an increase in oscillatory alpha (8-13 Hz) power, a well-established signature of neural inhibition [5,6], in occipital brain areas. During a later time window, 1500ms after the cue, intentional forgetting was associated with reduced employment of active rehearsal processes, as reflected by an attenuated upregulation of item-specific representations as compared to intentionally encoded items. Our data show that active inhibition and selective rehearsal are two separate mechanisms whose consecutive employment allows for a voluntary control of memory formation.

## Results & Discussion

### Forgetting instructions reduce subsequent memory performance

We employed the well-established item-method directed forgetting paradigm [1,2,4] in order to investigate the mechanisms underlying intentional forgetting (Figure 1A; see STAR Methods). As expected, TBF items were remembered worse than TBR items (Pr: hit rate - false alarm rate, M_TBF_=0.46, M_TBR_=0.56, t(17)=4.19, p=0.0006, Figure 1B). The reduced recognition memory performance suggests that the intention to forget or remember indeed leads to the down- or upregulation of individual memory traces [1,7].

**Figure 1:**
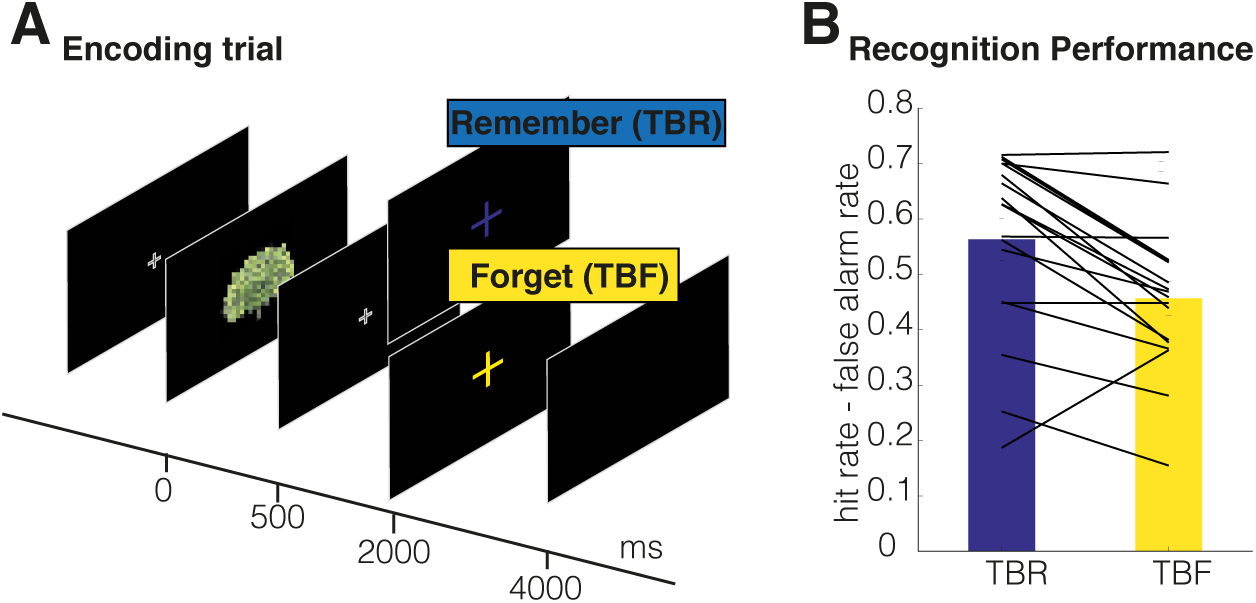
Paradigm and behavioral results. (A) Each encoding trial contains a picture of an everyday object, followed by either a TBR (to be remembered) or a TBF (to be forgotten) cue. (B) TBF cues led to reduced recognition memory performance relative to TBR cues in almost all participants; lines, performance of single participants. Detailed information on trialnumbers in each condition is provided in Figure S1.

### Item-cue similarity as a measure of active inhibition and active rehearsal

It is unclear whether reduced memory for TBF items is due to their active inhibition – i.e., a specific downregulation of their neural representations (Figure 2A, pink background) – or caused by reduced recruitment of rehearsal processes (Figure 2A, green background). On a behavioral level, both accounts predict reduced memory for TBF items and are therefore indistinguishable. However, the two accounts make dissociable and testable predictions regarding the fate of item-specific representations after a TBF and a TBR cue (Figure 2A).

**Figure 2:**
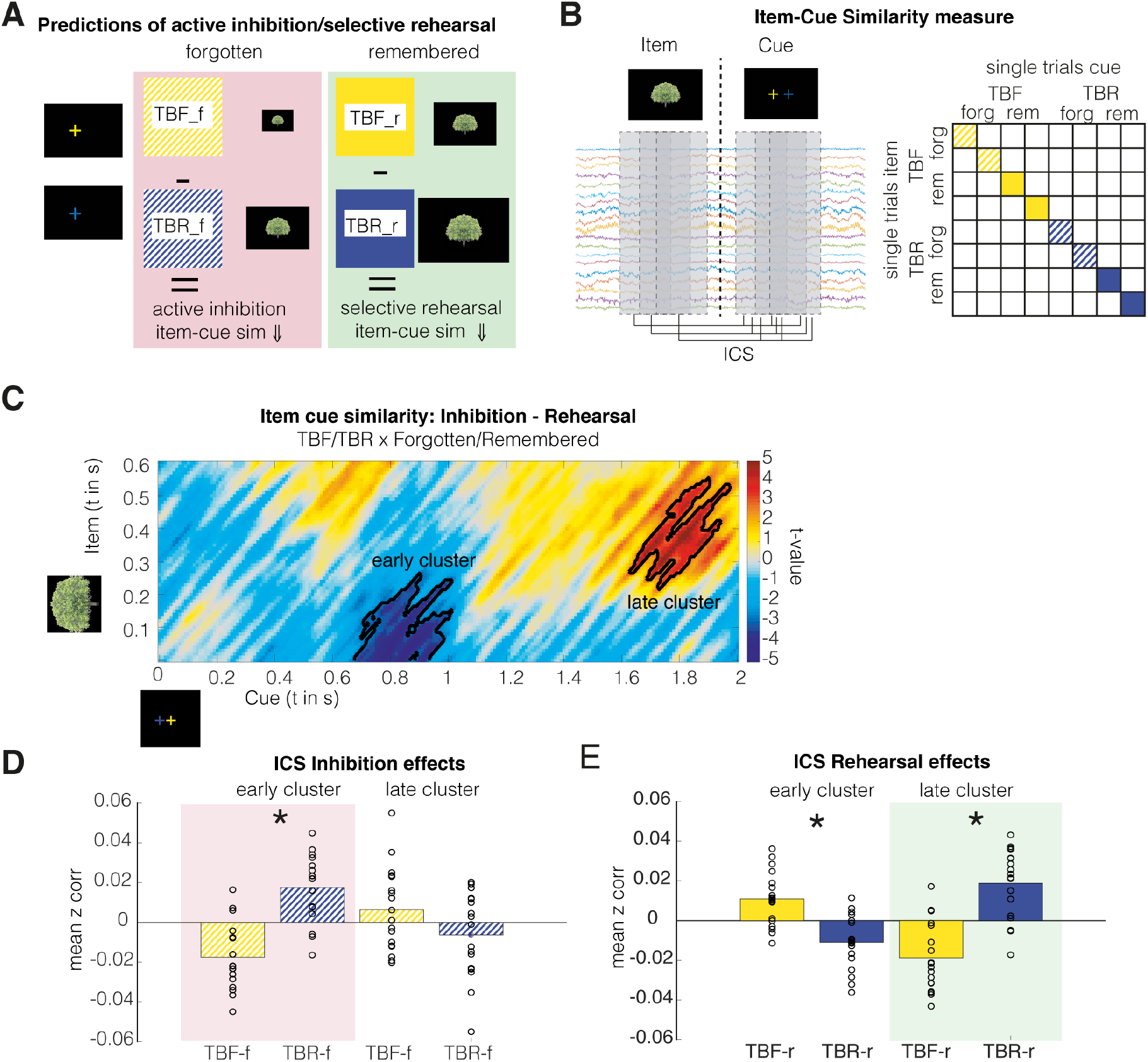
Transformation of item-specific representations during directed forgetting. (A) Predictions of active inhibition and selective rehearsal accounts on item-cue similarity (ICS) (B) Correlations between EEG activity during the item and the cue interval in every trial were calculated as single-trial ICS measures for all combinations of item-time x cue-time intervals. Trial-based ICS values were contrasted depending on the cue (TBF or TBR) and subsequent memory (remembered or forgotten). The matrix shown on the right depicts the correlations scheme included in the analysis. (C) Interaction effect (TBF/TBR x memory) testing for the differential employment of active inhibition (TBF-f vs. TBR-f) or reduced rehearsal (TBF-r vs. TBR-r). Colors depict t-values of ICS differences; significant clusters are highlighted by black contours (p_corr_<0.05). (D,E) Mean ICS values of forgotten (D) and remembered (E) items in the two interaction clusters shown in (C). The early negative cluster shows a significant inhibition effect, the late positive cluster shows significant reduced rehearsal effect. ICS values were mean centered to subject-specific mean ICS values for forgotten or remembered items to depict within-subject differences. *, p<.05. Predictions for active inhibition and selective rehearsal effects are highlighted in pink and green, respectively. Figure S2 depicts several control analyses demonstrating that here presented results are not confounded by item unspecific effects.

According to the active inhibition account, representations of forgotten TBF items (TBF-f) should be attenuated below the level occurring during passive forgetting, i.e., they should be reduced as compared to the representations of forgotten TBR items (TBR-f). This effect should be specific to the contrast of active versus passive forgotten items and not occur for later remembered TBF and TBR items (TBF-r and TBR-r, respectively). By contrast, the selective rehearsal account proposes that reduced memory for TBF items results passively from reduced rehearsal. Accordingly, item representations of subsequently remembered TBR items (TBR-r) should be actively upregulated as compared to unintentionally remembered TBF items (TBF-r). This rehearsal-related upregulation of item representations should drive remembering of the items and should thus not occur for later forgotten TBR and TBF items.

The two accounts thus make clearly distinguishable predictions for the neural representations of items in the different conditions: The active inhibition account predicts that representations of TBF-f items are suppressed below the level of TBR-f items (TBF-f < TBR-f), and that this suppression does not occur for TBF-r or TBR-r items. The reduced rehearsal account predicts decreased representations of TBF-r as compared to TBR-r items (TBF-r < TBR-r), an effect that again should be specific to remembered items and not occur for later forgotten items. In order to isolate these two effects, we thus tested the interaction between the two contrasts [8,9], i.e. (TBF-f vs. TBR-f) - (TBF-r vs. TBR-r). In this interaction contrast, active inhibition should result in lower RSA values, while active rehearsal results in higher values (Figure 2A&B).

### Item-cue similarities provide evidence for early active forgetting and later reduced rehearsal

As a measure of item representations after the cue, item-cue similarities (ICS) were estimated in each trial by calculating Spearman’s rank correlations between spatiotemporal EEG activity patterns [10–12] during item presentation and those following the TBF/TBR cue presentation (Figure 2B; STAR Methods). ICS values were calculated for all combinations of item and cue time windows, resulting in a time-resolved matrix of item representations during the cue interval. This analysis allows for a time-resolved assessment of the specific predictions of the active inhibition and the reduced rehearsal accounts.

Cluster-based permutation statistics applied on the ICS matrix revealed an early negative (p_corr_=0.004) and a late positive cluster (p_corr_=0.009, see Figure 2C). The early negative cluster corresponds to EEG activity around 600-1000ms after the onset of TBF/TBR cues and 0-300ms after item onset. Follow-up analyses of simple contrasts in this early cluster revealed a significant decrease of ICS for TBF-f vs. TBR-f items (t(17)=-4.52, p=0.0003, Figure 2D), i.e., item-related patterns were reduced during intentional as compared to unintentional forgetting. This effect was not observed when comparing later remembered TBF and TBR items (see below), supporting the active inhibition account.

The second significant cluster occurred later, 1500-2000ms after cue onset and 250-600ms after item onset. In this late cluster, ICS was significantly higher for TBR-r than TBF-r items (t(17)=-4.5941, p=0.0002, Figure 2E). Follow-up analyses did not reveal active inhibition in this time period (TBF-f vs. TBR-f; t(17)=1.287, p=0.22, Figure 2D). This result suggests that remembering of TBR items is driven by an upregulation of the corresponding memory traces, and that this process is less recruited for incidentally remembered TBF items.

Interestingly, an “inverted” rehearsal effect was present in the early cluster as well: we observed higher ICS for TBF-r items as compared to TBR-r items (t(17)=3.43, p=0.0032, Figure 2E). TBF-r items are remembered despite the forget instruction. This effect may reflect a paradoxical rebound due to a failed inhibition attempt [13,14]. Since this effect occurred early after the cue, prior to the voluntary rehearsal effect, it may reflect an automatic reactivation of a strong item representation acting beyond inhibitory reach [13,15,16]. Failed inhibition attempts have been shown to lead to paradoxical rebound and enhancement effects [13,14]. Importantly, the post-hoc tests show that both, the active inhibition effect and the paradoxical rehearsal effect, reach significance, i.e. the interaction contrast (Figure 2C) is driven by both effects.

The time-resolved ICS analysis revealed inhibition effects that preceded rehearsal effects. Additionally, the analysis also revealed differences in the inhibited and rehearsed item information: While inhibition specifically affected early item information (0-300ms, Figure 2C), rehearsal occurred only for item representations in later processing steps (250-600ms). This dissociation suggests that inhibition specifically targets lower-level perceptual representations, whereas rehearsal acts on higher-level semantic representations, in line with previous findings [17,18].

To exclude possible confounds in our assessment of ICS, additional control analyses showed that condition-specific differences in ICS were indeed driven by item-specific correlations between item and cue representations and do not reflect unspecific condition effects (Figure S2).

### Increased posterior alpha oscillations precede active inhibition of item-specific representations

Alpha oscillations (~8-13Hz) have been described as a signature of active inhibition of local information processing [6,19–21]. In order to reveal the mechanisms underlying the up- and downregulation of memory traces, we thus investigated brain oscillatory dynamics related to active inhibition and selective rehearsal. Again, we contrasted activity during TBF-f vs. TBR-r items as an index of active inhibition, and during TBF-r vs. TBR-r items reflecting selective rehearsal.

We conducted a three-dimensional cluster permutation statistic across frequencies, time points, and electrodes, contrasting TBF-f with TBR-f trials. This analysis revealed a significant increase of alpha power after the TBF cue (~8-13Hz, 100-1000ms, p_corr_=0.001, Figure 3A). This effect included a similar time window as the active inhibition effect on ICS (600-1000ms; see above) but showed a markedly earlier onset (starting at 100ms rather than 600ms post cue onset). We therefore investigated alpha power changes separately for the time windows preceding and during the ICS effect, by dividing the power effect time window into equal halves (0-500ms and 500-1000ms, respectively). We applied beamforming-based source estimation both to the early alpha power effect (preceding the active downregulation of item-specific memory traces) and to the late alpha power effect (during this downregulation). This analysis revealed a striking pattern: The early alpha power increase was localized to two clusters located in the occipital cortex (MNI peak −21/-71/-9, left lingual gyrus, t(17)=4.38, p_corr_=0.017) and in the right anterior medial temporal lobe (MNI peak 42/-10/-40, t(17)=4.43, right inferior temporal gyrus, p_corr_=0.034). This early occipital alpha power increase preceded the active downregulation of memory traces. It was specific to TBF trials that were forgotten, and did not occur for later remembered trials (Figure 3B; interaction analysis, i.e. contrast of active inhibition vs rehearsal, see STAR Methods: t(17)=3.05, p=0.007; follow-up pairwise tests: TBF-f vs TBR-f: t(17)=5.18, p=0.000076; TBF-r vs TBR-f: t(17)=0.83; p=0.22). In contrast, the later alpha power effect was localized to occipital areas extending to posterior temporal, parietal, and posterior midline areas (Figure 3C, MNI peak −9/-29/40, left middle cingulate cortex; t(17)=4.84, p_corr_=0.02). Analysis of this later time window showed an unspecific main effect of increased alpha power for TBF vs. TBR items independent of later memory (Figure 3C, interaction analysis p=0.01, t(17)=1.74, main effect of TBF vs TBR: t(17)=3.46, p=0.003; pairwise tests: TBF-f vs TBR-f: t(17)=4.82, p=0.0002; TBF-r vs TBR-f: t(17)=1.52, p=0.15). Figure S3A illustrates the temporal dynamics and frequency range of the inhibition-related power changes.

**Figure 3:**
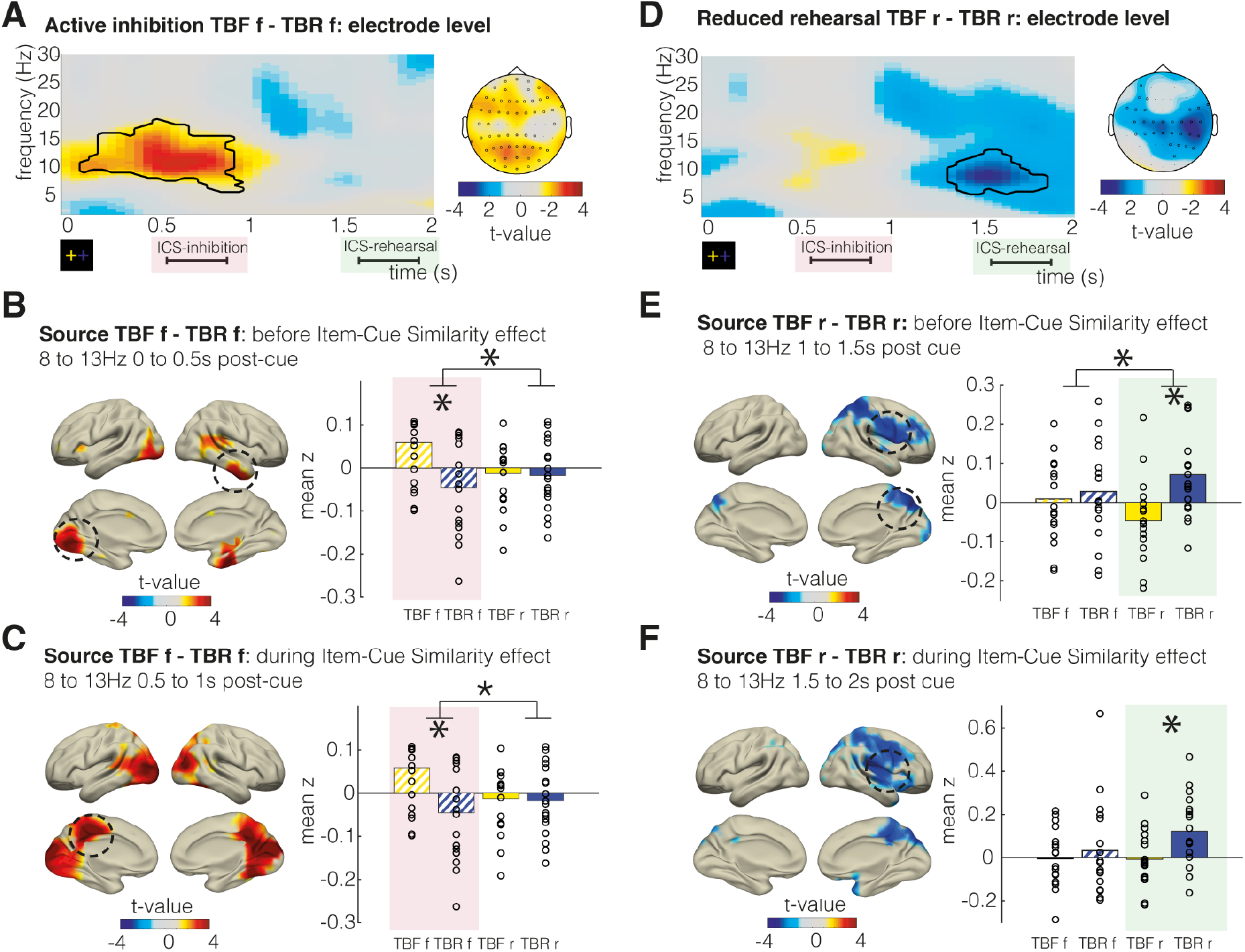
Brain oscillatory correlates of active inhibition and reduced rehearsal. (A) Contrast of intentionally forgotten trials (TBF-f) vs. incidentally forgotten trials (TBR-f). Results are corrected for multiple comparisons using a cluster-based permutation test across all electrodes, frequencies (2-30Hz), and time points (0-2s post cue). Time-frequency plots show mean t-values across all significant electrodes. Right: topographical plot highlighting the respective significant electrodes. (B) Oscillatory active inhibition effects preceding active inhibition-related ICS changes (time window: 0-0.5s post-cue). (C) Oscillatory active inhibition effects during the inhibition-related ICS changes (time window: 0.5-1s post-cue). (D) Significant differences of oscillatory power between incidentally remembered (TBF-r) and actively rehearsed trials (TBR-r), cluster permutation results as in (A). Please note that the selective rehearsal contrast is always plotted as reduced rehearsal (TBF-r – TBR-r). (E) Reduced oscillatory rehearsal effect preceding rehearsal-related ICS changes (time window: 1-1.5s post-cue). (F) Reduced oscillatory rehearsal effects concurrent with the rehearsal-related ICS changes (time window: 1.5-2s post-cue). Source plots show differences between TBF-r and TBR-r, thresholded at p<0.01, circles highlight significant clusters (p<0.05). Bar plots to the right depict average power in the significant source clusters, black stars indicate p<0.05, black circles denote single subject trial averages. Figure S3 additional shows timefrequency changes in the respective significant source clusters and additional analysis depicting the conjunction of inhibition and rehearsal effects.

Taken together, these results show early alpha power increases in occipital areas during the successful suppression of perceptual memory traces, reflected in the inhibition of early, perceptual parts of the item representation (Figure 2C) [22–24].

### Late alpha/beta power increases index selective rehearsal

Interestingly, alpha power increases did not only precede and accompany active inhibition, but also selective rehearsal. Alpha power increased for TBR-r vs. TBF-r items in a cluster that again preceded and concurred with the time period of the ICS selective rehearsal effect (3-dimensional cluster statistics; 8-13Hz, 1300-1700ms post-cue, p_corr_=0.03; Figure 3D). Again, we separately analyzed effects in the two time windows (1000-1500ms and 1500-2000ms post cue). In both time windows, rehearsal-related alpha power changes were localized to right occipital, parietal and frontal areas, preceding the ICS effect (two clusters, cluster 1: MNI peak 12/-70/39, right cuneus, t(17)=-4.55, p_corr_=0.018, cluster 2: MNI peak 64/-9/38-, right postcentral gyrus, t(17)=-3.75, p_corr_=0.019) and concurrent to the ICS effect (MNI peak 32/19/19, right insula, t(17)=-3.95, p_corr_=0.044). In the time window preceding the rehearsal ICS effect, a rehearsal-specific alpha effect was evident with stronger alpha power increases for TBR-r than TBF-f items (interaction: t(17)=2.40, p=0.029, follow-up pairwise tests: TBF-f vs TBR-f: t(17)=-0.55, p=0.59 TBF-r vs TBR-f: t(17)=-4.41; p=0.0004). Concurrent to the ICS effect, alpha power changes were not rehearsal-specific, but related to a general difference between the TBF and TBR condition (interaction: t(17)=1.40, p_corr_=0.18, TBF vs TBR t(17)=-2.66, p_corr_=0.017). Figure S3B further illustrates these effects, showing the time-frequency resolved power changes in the respective source clusters.

The relative increases in alpha power for successfully rehearsed TBR-r items are well in line with previously described increases in alpha power during working memory maintenance [25]. Interestingly, while the oscillatory power changes related to active inhibition were restricted to the alpha frequency range, the selective rehearsal effect extended to higher frequencies in the beta range (Figure S3A&B), possibly reflecting maintenance processes [26,27].

Taken together, we found robust increases in alpha power related to both active inhibition and selective rehearsal that clearly precede the onset of ICS effects. We propose that alpha power increases during active inhibition reflect an active downregulation of unwanted memory traces, whereas active maintenance during selective rehearsal indicates a focus on internal representations which also requires a suppression of bottom-up information flow of possibly distracting information [6,28,29]. This interpretation is also in line with the different sources of alpha power changes: While the early alpha inhibition effect was localized to occipital regions and thus may indicate a downregulation of visual representations, the alpha power increases during selective rehearsal were found in right-lateralized parietal regions and possibly index task demands [30].

### Alpha power changes are spatially overlapping to ICS changes

In order to investigate the spatial extent of ICS effects and to evaluate the overlap of ICS and alpha effects, we conducted an ICS searchlight analysis. ICS values were calculated for every source voxel and its 63 nearest neighbors employing the same sliding window spatiotemporal ICS calculation approach as on the sensor level (see Figure 2B & Figure 4). The searchlight analysis focused on the two significant temporal clusters reported in Figure 2C. This analysis revealed that ICS inhibition effects (0-300ms item time, 600-100ms cue time) were strongest in occipital areas (Figure 4A), ICS rehearsal effects (270-600ms item time, 1600-2000ms cue time) were most pronounced in parietal areas (Figure 4C).

**Figure 4:**
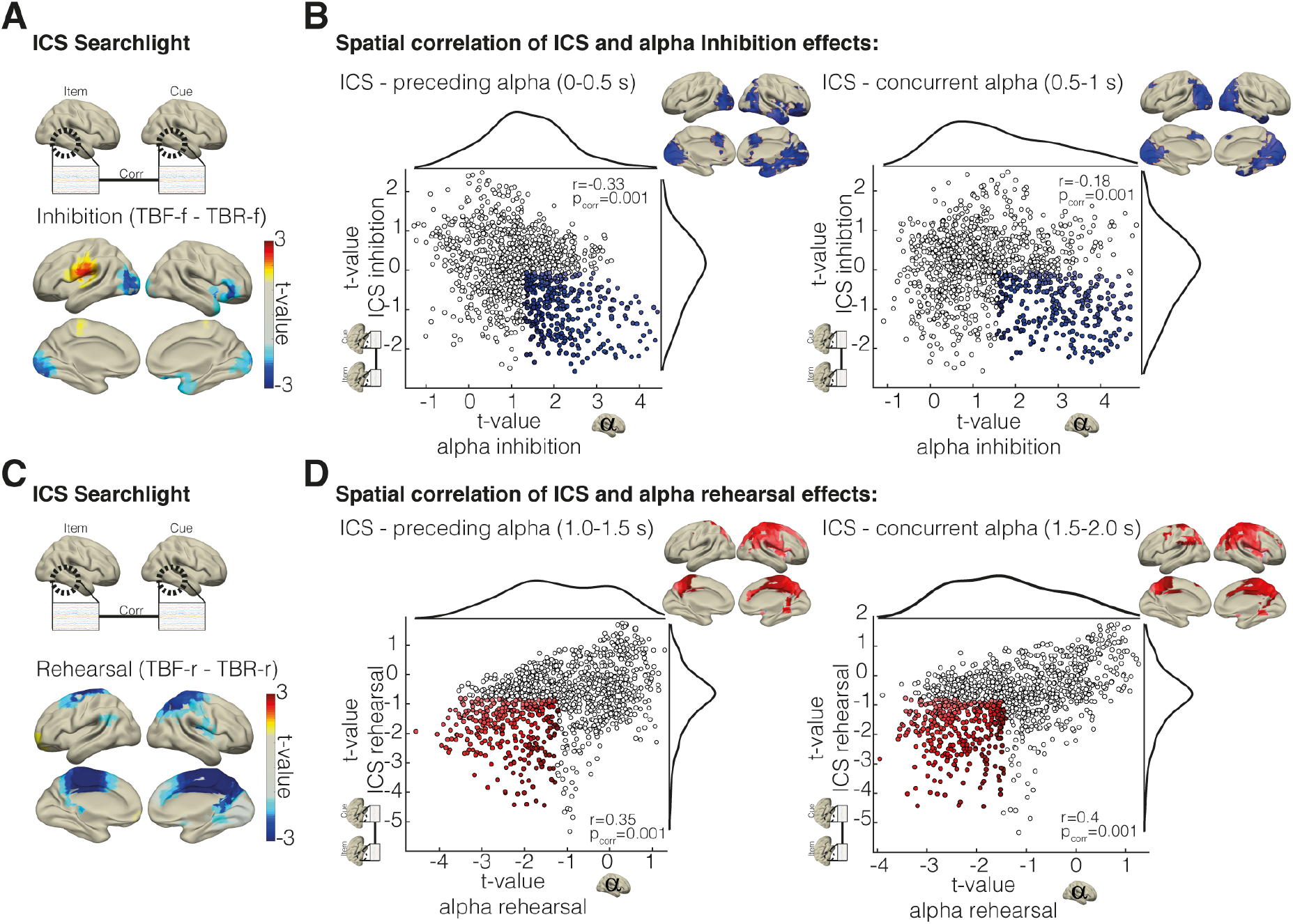
ICS searchlight analysis and overlap with alpha power effects. (A) Results of searchlight ICS analysis, t-maps of inhibition and (C) rehearsal ICS effects, (threshold p<0.05 uncorrected). (B&D) Correlation of alpha power effects and ICS searchlight effects. Inhibition (B) and rehearsal (D) alpha and ICS searchlight effects were correlated across voxels. Scatterplots depict single voxels. To illustrate the spatial overlap, voxels exhibiting overlapping effects, highlighted in blue and red, were mapped back onto the brain surface. Highlighted voxels were selected using the respective mean t-values for alpha and ICS effects as a cutoff (i.e. decreased ICS & alpha power increases for inhibition effects). Figure S4 shows the correlation of forgetting success to the spatial overlap of alpha power and ICS inhibition effects.

To evaluate the spatial overlap of ICS and alpha power effects, we correlated the topographical patterns of ICS and alpha effects across all voxels. This analysis revealed significant negative correlations between ICS inhibition effects and preceding and concurrent alpha power inhibition effects (Figure 3B). This relationship was stronger for alpha power effects preceding ICS inhibition effects than for alpha power changes concurrent to ICS effects (Pearson and Filon’s z, z=-5.1460, p <0.001, [31]). Rehearsal effects were positively correlated: Areas exhibiting stronger alpha power increases for TBR-r vs TBF-r trials showed stronger ICS increases (Figure 4D). This relationship was stronger for concurrent than preceding alpha power changes (Pearson and Filon’s z, z=-5.9458, p<0.001). These results suggest that alpha power changes up- and downregulate regionally specific item representations.

Interestingly, specifically alpha power changes preceding ICS inhibition effects correlated strongly to ICS effects, suggesting that the spatial overlap of alpha power changes and item representations might predict forgetting. Thus, we correlated the spatial overlap of early alpha power increases and ICS effects with behavioral forgetting success. The ICS-alpha overlap was evaluated by correlating mean alpha power maps (0-500ms post cue) to mean searchlight ICS maps (0-300ms item and 600-1000ms cue time window), in every condition and subject. A significant correlation was found only for TBF-f items, where a higher overlap of alpha power and ICS was related to stronger forgetting effects across participants (r=0.52, p=0.027, Figure S4). This suggests that the spatial precision of alpha power changes targeting item representations predicts successful voluntary forgetting.

### Active inhibition and selective rehearsal are dissociable processes contributing to directed forgetting

Inhibition and rehearsal effects showed no temporal overlap, neither considering alpha power effects nor ICS effects (see Figure S4C&D for a conjunction of effects). Our data provides evidence that directed forgetting relies on two temporally separable employed processes, active inhibition and selective rehearsal. To confirm this, we investigated whether participants showing stronger active inhibition effects also exhibit more pronounced rehearsal effects. Correlating inhibition and rehearsal effects across subjects separately for ICS and alpha power effects (extracted from significant source clusters) showed no significant correlation for either of these variables (inhibition & rehearsal ICS effects: r_spearman_=-0.23, p=0.34; early alpha power inhibition & rehearsal effects: r_spearman_=-0.14, p=0.55; late alpha power inhibition & rehearsal effects: r_spearman_=-0.02, p=0.95). This result suggests that active inhibition and selective rehearsal are independently employed [32].

### Intentional forgetting relies on flexible modulation of memory representations

The present results show that intentional forgetting of memories relies on two different processes, an active downregulation of unwanted memory traces followed by a reduced rehearsal of these traces. Interestingly, a prior directed forgetting intracranial EEG study reported similar increases in alpha power together with local theta oscillations involved in downregulating hippocampal activity around the time window of our current inhibition-related ICS and alpha power effects [8]. Going beyond oscillatory correlates, our results demonstrate that item representations are indeed actively inhibited at the time of these hippocampal effects. Together, these findings show that motivated forgetting attenuates memory traces, leading to their later forgetting.

## Acknowledgments

This research was funded by the Deutsche Forschungsgemeinschaft (DFG, German Research Foundation)—Projektnummer 316803389—SFB 1280, as well as via Projektnummer 122679504 – SFB 874 (awarded to NA). We thank Malte Kobelt for assistance with data acquisition and preprocessing.

## Author Contributions

M.C.F & G.T.W conceived, designed and executed the experiment, M.C.F wrote analysis scripts, analyzed the data and wrote the initial draft, G.T.W, N.A. provided expertise and feedback and reviewed and edited the draft, N.A. acquired the funding for the research.

## Declaration of Interest

The authors declare no competing interests.

## METHODS

### RESOURCE AVAILABILITY

#### Lead Contact

Further information and requests for resources should be directed to and will be fulfilled by the Lead contact, Marie-Christin Fellner.

#### Materials Availability

This study did not generate new materials.

#### Data and Code Availability Statement Examples

Data and scripts to carry out all reported analysis are publicly available (DOI 10.17605/OSF.IO/UPKWE, osf.io/upkwe)

### SUBJECT DETAILS

#### Participants

Twenty-three healthy volunteers with normal or corrected-to-normal vision took part in the experiment. One dataset was excluded because of the age of the participant (48 years), another 4 datasets were excluded after artifact correction (less than 10 trials in one of the conditions), resulting in a final sample of 18 datasets (mean age: 23.4, age range: 18-32 years, 8 male). All reported analysis is based on this sample of 18 datasets. All subjects were right handed and reported no history of a neurologic or psychiatric disease. All participants gave their written informed consent, and the experimental protocol was approved by the ethical review board of the Faculty of Psychology at Ruhr University Bochum.

### METHOD DETAILS

#### Experimental design

The experiment consisted of three parts: an encoding phase, a short distractor task (3min of backwards counting starting from random 3-digit numbers), and a recognition phase. During the paradigm, participants were seated in front of a computer screen. The experiment started with the instruction on screen and 4 practice trials. Participants were explicitly instructed that the experiment served to study the effect of forgetting on memory. Pictures of everyday objects were presented, followed by fixation cross cueing whether the preceding item was to be forgotten (TBF) or to be remembered (TBR). Depending on the color of the fixation cross, the participants were instructed to either rehearse and try to encode the previously presented item, or to voluntarily forget the item as these items would not be tested later. Participants were instructed not to remember the TBF items in order to improve remembering the crucial TBR items. After a brief distractor phase, they performed an item recognition task which nevertheless contained all TBF and TBR items that had been presented during encoding, randomly intermixed with new pictures. Participants were explicitly asked to remember both, TBF and TBR items.

We used 288 picture stimuli depicting nameable objects from an existing database [33]. During encoding, 96 pictures were presented and followed by a TBF or TBR cue, respectively. The sequence of TBF and TBR trials was randomized. Each trial started with a fixation cross (500-1000ms jittered), then the picture was presented for 500ms, followed by another fixation cross (1500ms) and the TBF/TBR cue that remained on screen for 2000ms. During recognition, all old items (TBF&TBR) were shown mixed with 96 new items. Participants were instructed to respond to each item on a 1-6 confidence scale (i.e. 1= very sure old, 6 = very sure new). Participants used index, middle and ring fingers on both hands to respond. Each recognition trial started with a fixation cross (500-1000ms jittered), then the item was presented for 500ms, followed by another fixation cross (1500ms) and a response screen for 2000ms. During all phases, a blank screen was presented during the inter-trial interval (1250ms). Picture material, response hand and TBR/TBF cue colors were counterbalanced across the participant sample.

#### EEG recording and preprocessing

EEG was recorded from 64 electrodes in an extended 10/20 montage (BrainAmp Standard, EasyCap). Recordings were referenced to Fz and later re-referenced offline to average reference. Impedances were kept at below 10kΩ. The signals were amplified between 0.1Hz and 250Hz and recorded with a sampling rate of 500Hz.

All EEG data preprocessing and analyses were carried out using fieldtrip (http://www.fieldtriptoolbox.org, [34]) and custom MATLAB scripts. All scripts and preprocessed data are available (osf.io/upkwe). Data were epoched in trials from 1 second before an item to 5 seconds after item onset during encoding. Data were visually inspected to exclude trials with idiosyncratic artifacts (channel jumps, muscle artifacts, noisy channels) from further analysis. Noisy channels were excluded (in four datasets, up to three electrodes were excluded). Infomax independent component (IC) analysis was applied to correct for residual artifacts (e.g., eye blinks, eye movements, or tonic muscle activity). On average 29.39 TBF-f (range: 11-52), 52.61 TBF-r (range: 34-66), 20.11 TBR-f (range: 10-52), and 63.11 TBR-r trials (range: 30-83) passed artifact corrections. A more in-depth summary of trial numbers and the impact of trial sample size on alpha power estimates is shown in Figure S1.

#### Representational similarity analysis of item-cue similarity

Item-cue similarities were calculated by correlating EEG activity during item presentation (0-500ms) with EEG activity post-cue presentation (2000-4000ms relative to item, 0-2000ms post-cue) within each trial. To this end, artifact-corrected raw data was downsampled to 100Hz. To remove mean ERP-related activity, all trials were normalized using the mean and standard deviation across all trials for each data point and channel. We ensured that our normalization procedure did not induce any spurious effects. As we applied a z-transformation to the EEG activity at every time point across all trials of all conditions in order to remove ERP components prior to calculating correlations, we also ran a control analysis, calculating item-cue similarity using data that were not z-transformed across conditions but only within conditions (Figure S2A). This approach yielded the same pattern of results.

EEG data was cut into windows of 200ms in overlapping time windows with 10ms increment, yielding a matrix of 21 time points and 64 electrodes every 10ms. These two-dimensional matrices were then concatenated into one-dimensional vectors with combined spatial and temporal information (vector dimensions: 21 time points x 64 electrodes). Spearman correlations were calculated for every combination of time x channel vectors during item representation and time x channel vectors during the cue period. This results in item-cue similarity matrices for all combinations of item time bins and cue time bins.

#### Searchlight ICS analysis

To investigate spatially resolved ICS effects, ICS was calculated for searchlights centered around every virtual source electrode. The size of the searchlight was set to 64 voxels (corresponding to ~4.5cm diameter). As source projected EEG still has a low spatial resolution (here 1cm^3^ virtual electrode spacing), only relatively few fully spherical searchlights can be “moved” across the brain, leading to a bias to midline voxels. Therefore, searchlights were defined not as spheres but as the 64 voxels nearest to the selected voxel including itself.

Using these searchlights consisting of 64 virtual source electrodes, ICS was calculated in the item-cue time windows that were identified in the sensor-level analysis: An early cluster between 0-300ms with regard to item onset and 600-100ms after the cue; and a late cluster between 270-600ms after the item and 1600-2000ms after the cue (see Figure 2C, selected windows correspond to highlighted clusters). In these item x cue time windows, ICS was calculated for every virtual source searchlight using the same sliding window approach as for the sensor level data (200ms of data concatenated across all included electrodes). The resulting correlation values were averaged across the selected time windows, resulting in one average ICS value for each virtual source searchlight, condition, and subject. T-contrasts were calculated to explore the effects of interest (inhibition contrast and selective rehearsal contrast). Results of these contrasts were mapped onto a brain surface model (Figure 4).

#### EEG oscillatory power analysis

Data were filtered using wavelets with a length of 5 cycles to obtain oscillatory power between 2 and 30Hz. Single trial power values were z-transformed using frequency- and channel-specific means and standard deviations across all trials.

Source analysis was performed using a linearly constrained minimal variance (LCMV) beamformer [35], calculating a spatial filter based on the whole length of all trials. For all subjects, we used a standard boundary element source model with a grid resolution of 10mm based on the Montreal Neurological Institute (MNI) brain and standard electrode positions realigned to the MNI MRI. The source timecourse for each grid point was calculated, subjected to a wavelet analysis (same settings as for the raw data) and z-transformed as for the electrode-level data. Statistical analysis was restricted to grid voxels inside AAL-defined brain regions. For region of interest analyses, data across all grid voxels covering the region of interest (significant source) were averaged. Grid voxel data were interpolated to a 2mm resolution single-subject MNI brain for plotting and to define the locations of clusters and peaks.

### QUANTIFICATION AND STATISTICAL ANALYSIS

#### Behavioral Analysis

To analyze memory performance, we used a signal detection approach to obtain bias free measures of memory strength and to classify hits and misses relative to an individually defined neutral response criterion (for a similar procedure see [36,37]). As demonstrated previously, individually defining the confidence rating boundary between hits and misses enhances signal to noise ratio by taking into account individual differences in the use of confidence ratings [36].

First, the 1-6 confidence recognition responses of the participants were mapped to binary old/new judgements taking into account participants’ individual response biases. Confidence ratings were defined as “old” judgements up to the highest rating that was more often used for new items than old items during recognition. According to this definition, for two participants only “1” indicated an old response, for 4 participants “1-2”, for 9 participants “1-3” and for 3 participants “1-4”. Memory performance was then assessed by calculating Pr, i.e. the difference between hit rate (percentage of old items remembered as old) and false alarms (percentage of new items incorrectly judged as old).

#### Statistical analysis ICS

For statistical testing of significant ICS differences, a cluster-based permutation approach was used to accommodate for potential biases due to different trial numbers in the different conditions and to correct for multiple comparisons. To this end, the trial labels were shuffled in each subject 1000 times. This random data was then used to construct null distributions of effects under the existing bias in trial number. In a second, group statistical step, clusters of temporally adjacent significant differences (threshold p<0.01) were identified and the sum of t-values in each cluster was calculated in the original data and in the data based on the random permutations. If no t-value reached significance in one of the permutations, a cluster value of 0 was assigned. Significance of clusters was assessed by calculating the rank of the cluster t-values in the distribution of random data (reported as p_corr_). A cluster was interpreted as significant if an absolutely higher cluster t-value was found in less than 5% of the random permutations.

#### ICS control analysis

To ensure that the ICS effects indeed reflect reactivation of item-specific information, we conducted several control analyses. A possible confound of the reported ICS interaction effects reported in Figure 2C is that effects might not only reflect modulations of item-specific representations, but instead general condition-specific activity differences such as ERPs, power changes, or other unspecific differences.

A possible confound is whether the reported effects – even if they rely on single trials rather than trial-averaged ERPs – are indeed attributable to item-specific representations or whether they reflect differences that are independent on item identity. If the reported ICS effects are indeed attributable to item-specific memory traces, these effects should be specific to correlations of activity during one item window and activity during the matching cue window (i.e., within-item correlation); they should not be present when correlating activity during one item window with activity during a non-matching cue window (i.e., between-item correlation; see model matrix in Figure S2B).

Item specificity is often ensured by calculating the difference of within-item correlations and between-item correlations [38,39]. In Figure S2B we report an ICS analysis based on within-item – between item contrasts. We replicated the ICS analysis as shown in Figure 2 by contrasting first within-item ICS to between-item ICS (ICS_within-between_) in each condition. The interaction contrast presented in Figure S2B hence shows (TBF-f_within-between_ – TBR-f_within-between_) vs. (TBF-r_within-between_ – TBR-r_within-between_). In order to assess statistical significance of the contrasts, again a cluster-based permutation statistics was applied, randomly shuffling the assignment of single trials in each subject. This analysis revealed the same results as the main analysis: an early negative cluster around 600-1000ms post cue (p_corr_=0.001) and a late positive cluster around 1600-2000ms (p_corr_=0.024).

As within-item correlations (i.e., matching item and cue) in our data are necessarily within-trial correlations, and between-item correlations are across trials, directly calculating this difference may lead to biased results, as differences may not only be driven by within-between item effects but also by within-between trial effects. We therefore additionally tested whether similar effects as observed within-items occurred also between-items. We repeated the same analysis as reported in Figure 2C based on between-item correlations and tested for the reported interaction effects. Correlations of non-matching item-cue intervals were calculated and again the interaction between TBF/TBR x memory was assessed.

This analysis yielded a very different pattern of results than the within-trial item-cue similarity analysis (Figure S2C). Most importantly, the results are non-overlapping with the reported interaction effect (Figure 2C). Early after the cue, a significant positive effect was evident, an effect that is in stark contrast to the reported negative cluster in the within-item ICS analysis. This control analysis provides further evidence that the reported effects are indeed attributable to an up- or downregulation of item-specific representations.

Additionally, to ensure that the employed spatiotemporal correlation approach is sensitive to item specific effects, we calculated item-specific encoding-recognition similarity (ERS). Analyzing ERS allows one to assess item-specificity without the within-item/within-trial confound that biases ICS item-specific contrasts. ERS was calculated using the same spatiotemporal sliding window approach and trial permutation based cluster statistics as used in the main analysis. ERS was calculated for every condition separately. To reveal item-specific effects, within-item ERS was contrasted to between-item ERS in every condition. This analysis revealed for TBF-r, TBR-r, TBF-f significant item-specific clusters and a trend of an item-specific effect in the TBR-f condition (see Figure S2D).

Furthermore, we tested for a possible influence of ERPs on the reported ICS effects. ERPs were previously reported to show condition differences during directed forgetting [40]. ERP rehearsal, inhibition and interaction effects were evaluated and are reported in Figure S2E. These differences in average condition-specific waveforms reflecting attentional and mnemonic processes could potentially influence the ICS measure, which could substantially change the interpretation of our effects: ICS effects would then reflect condition-specific differences in neural activity and not up- or downregulations of item-specific memory traces. However, additional analyses demonstrated that ERP effects cannot explain the item-cue similarity pattern that we had observed: Conducting the same interaction analysis as reported in Figure 2C using trial-averaged ERPs rather than single-trial data did not yield any significant clusters. For this analysis single subject, condition specific ERPs were calculated and subjected to the same item-cue-correlation analysis (Figure S2F). More importantly, the pattern of ERP similarity values showed no comparable increases and decreases as observed during the ICS interaction analysis. The pattern of (non-significant) effects is actually opposing the ICS effects: numerically positive differences were found early after the cue followed by numerically negative differences late post-cue (Figure S2F). This control analysis shows that the ICS effects cannot be attributed to univariate condition differences.

Finally, we investigated whether our ICS results were directly driven by the reported differences in alpha/beta power between conditions (see main Figure 3). We thus repeated our main analysis of ICS after applying a bandstop-filter between 7-20Hz. This analysis yielded very similar results as our main analysis (Figure S2G), with the exception of lacking aliasing effects that are caused by (usually prominent) alpha oscillations.

#### Statistical analysis oscillatory power

For statistical analysis of EEG data, power spectra for each subject were collapsed and averaged across all trials for each “cell” of the design matrix (TBF-f, TBF-r, TBR-f, TBR-r). All of the following EEG analyses were based on these first-level averages (4 cells, 18 subjects). This prior averaging of data within each cell for each subject controls for possible biases of trial numbers in the analysis of main effects (e.g., there were more TBF-f trials than TBR-f trials, and thus, if all TBR trials irrespective of memory would be pooled, condition effects would be confounded by subsequent memory differences, see Figure S1A). This analysis is essentially equivalent to a 2×2 repeated measurements ANOVA.

For statistical analysis, we again used nonparametric cluster-based permutation tests as implemented in fieldtrip [41]. The cluster-based permutation test consisted of the following two steps: first, clusters of coherent t-values exceeding a certain threshold (here, p<0.01) along selected dimensions (time, frequency, electrodes/grid voxels) were detected in the data. Second, summed t-values of these clusters were compared to a null distribution of t-sums of random clusters obtained by permuting condition labels across subjects. This procedure effectively controls for type I errors due to multiple testing. The clusters of t-values subjected to permutation testing can be built across different dimensions: clustering can be performed on non-averaged data across all dimensions (electrode, frequency, time) or in a specific dimension when averaging over certain dimensions (i.e., averaging in the time-frequency window and then clustering across the electrode dimension). Clustering was employed along different dimensions depending on the data. In order to investigate power changes on the source level, power values of significant source clusters were extracted, averaged across significant voxels, and subjected to further statistical analysis. In order to correlate the active inhibition and selective rehearsal power effects across subjects, average power in the reported peak voxel of the effect was extracted.

To further ensure that reported significant alpha power changes are not confounded by possible biases cause by differences trial numbers between conditions, reported alpha power results were follow up by an additional trial based cluster permutation statistic clustering across sensors, similar to the reported ICS statistical analysis. Here, instead of shuffling the condition label of mean power values in each subject, trial labels in each subject were shuffled to construct a null distribution of the selected contrast (see Figure S1C). Importantly, this analysis yielded very similar results to the condition shuffling approach, demonstrating that reported results are not caused by trial number biases.

## Supplemental Material

**Figure S1:**
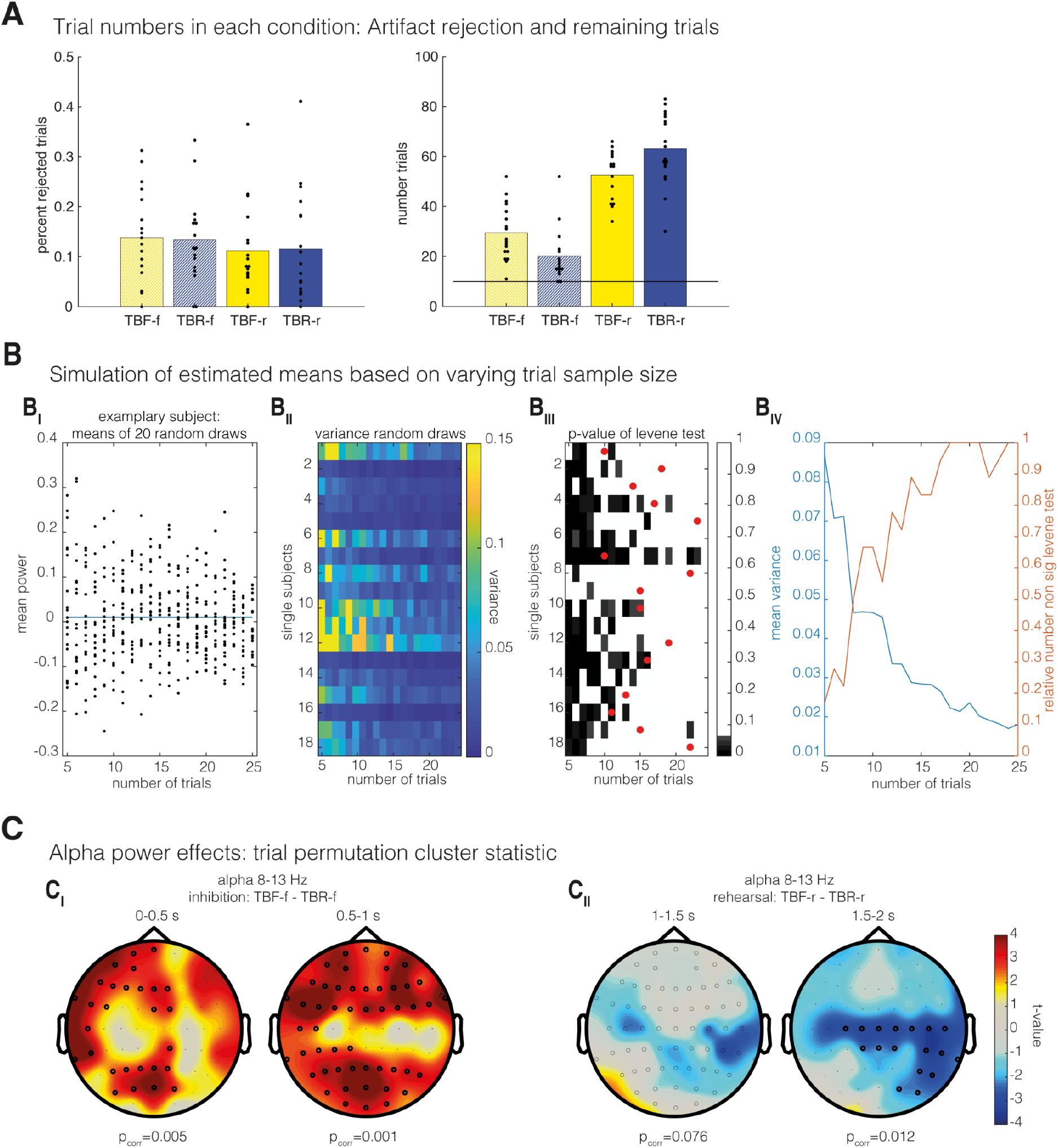
Analysis of trial number effects related to Figure 1. (A) Left: Relative trial numbers rejected during preprocessing in each condition. Right: Absolute trial numbers after artifact correction (subject inclusion criteria of 10 trials minimum highlighted by horizontal line). (B) Influence of trial numbers on the estimation of mean alpha power. A simulation was run randomly drawing samples of 5 to 25 trials, thereby generating 20 random samples for each trial number size and calculating the mean alpha power (8-13Hz, 0-2s past cue, POz) in each sample. (B_I_) Simulated means in one exemplary subject. Dots depict means, the blue line shows mean across all existing trials (n=174) in one subject. The variance between the black dots is clearly decreasing with more trials. (B_II_) Variances of simulated means for all subjects and trial numbers. (B_III_) P-values of Levene’s test of variance homogeneity, comparing the variance across multiple random drawings in differently sized subsamples to a sample of 25 trials (uncorrected for multiple comparisons). Red dots highlight the number of trials in the condition with the smallest number of trials in each subject. (No dots for subjects with more than 25 trials). This analysis clearly confirms that for the vast majority of subjects the available trial sample size is above the number needed for stable mean estimation. (B_IV_) Mean variance for each trial sample size and relative number of subjects showing no significant differences in variances for the different sample sizes. This plot indicates that variances are relatively stable in samples of 10 or more trials. (C) To further establish that alpha power results were not biased by differences in trial numbers between conditions, a trial-based permutation statistic of alpha power effects was carried out. Trial-based permutation statistics (as used for the ICS analysis, Figure 2) controls for potential trial biases by assessing statistical significance using null distributions based on the original data distribution. Trial labels of the tested contrasts were randomly shuffled in each subject. Using this trial-based permutation approach, we tested for (C_I_) alpha power inhibition effects preceding (0-500ms) and concurrent (500-1000ms) to ICS inhibition effects, and (C_II_) alpha power rehearsal effects preceding (1000-1500ms) and concurrent (1500-2000ms) to ICS rehearsal effects. This analysis largely replicated the results of the main analysis of inhibition and rehearsal effects on the sensor level (Figure 3), showing that alpha power effects remain stable also when controlling for trial number biases.

**Figure S2:**
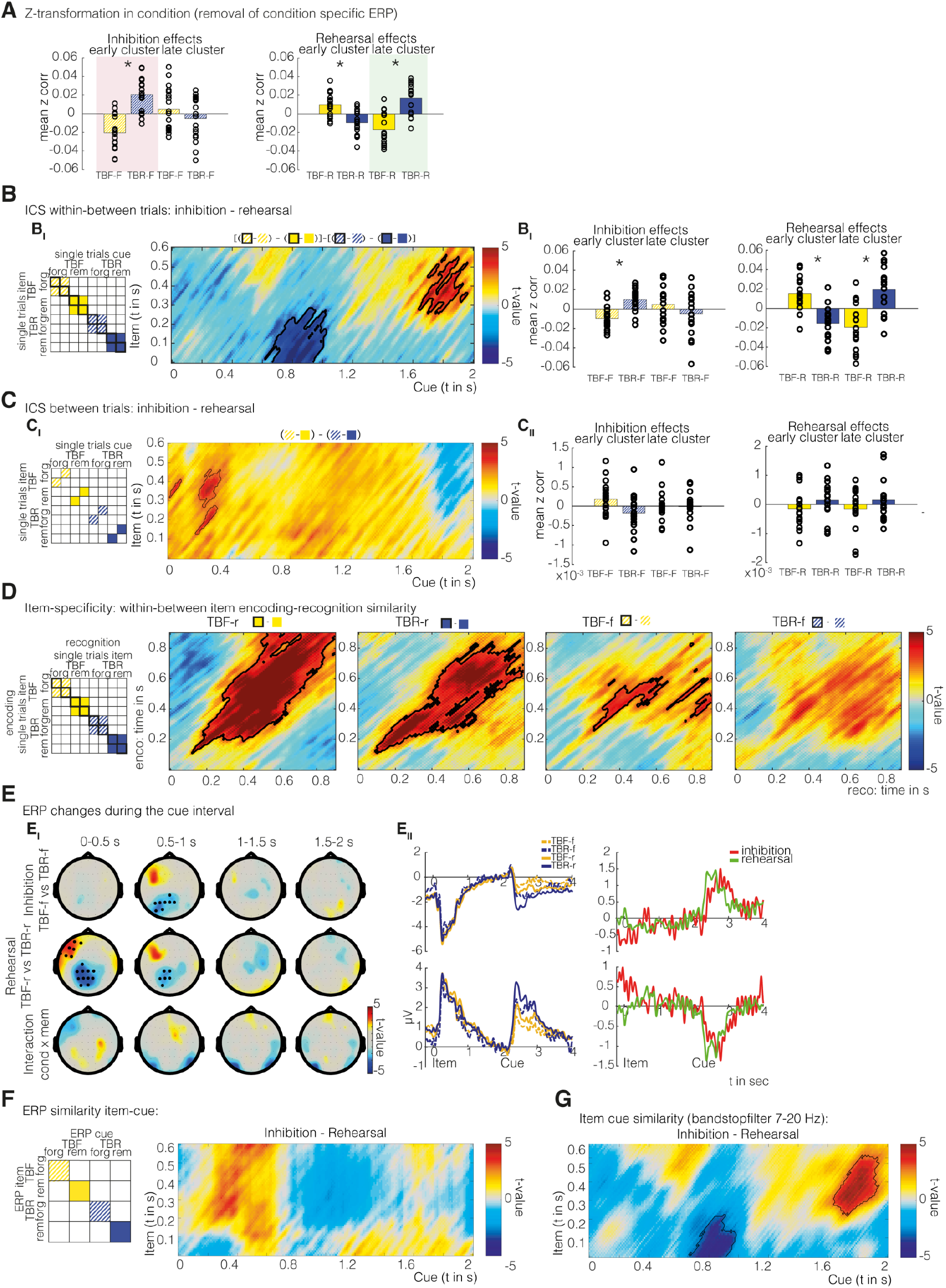
Item-cue similarity control analysis related to Figure 2. To ensure that item-cue similarity differences presented in Figure 2C were not confounded by item-unspecific differences between conditions, several control analyses were carried out. (A) Control analysis for normalization procedure. To ensure that different normalization steps prior to correlations did not influence our results, the same analysis as presented in Figure 2 D&E was reproduced employing condition-specific normalization prior to the item-cue similarity analysis. (B) Interaction effect based on within-item vs between-item ICS analyses. (B_I_) A cluster permutation statistic shuffling single trial labels revealed very similar results to the with-item results (compared to Figure 2 in the main manuscript, negative cluster p_corr_=0.024, positive cluster p_corr_=0.001). Contours highlight significant clusters. (B_II_) Mean ICS values based on within-item vs between-item ICS separately for forgotten and remembered trials in the two interaction clusters highlighted above. Note that the pattern of results remains unchanged. (C) Interaction effect based on between-item ICS analyses. (C_I_) The same analysis as in Figure 2C was also conducted between item and cue periods of different trials, revealing a very different result than within item analyses. (C_II_) Mean ICS values of between-item ICS in the interaction clusters highlighted in B_I_, note that no significant inhibition or rehearsal effects were present in the between-item ICS, demonstrating that reported ICS effects are specific to the within-item contrasts. (D) Item-specific encoding-recognition similarities in each condition. Outlined contours show significant clusters of item-specific information obtained by a cluster permutation statistic. Item-specificity was tested by permuting item labels in each subject and for each condition, providing a null distribution of no item-specific effects (TBF-r p_corr_=0.001, TBR-r p_corr_=0.001, TBF-f p_corr_=0.01, TBR-f p_corr_=0.072). (E) Condition-specific ERP effects. (E_I_) Topographical plots of inhibition, rehearsal and interaction effects in 500ms windows post TBR/TBF cues (cluster permutation tests across electrodes). Highlighted black dots show clusters of significant electrodes (p_corr_–0.05). (E_II_) Condition-specific ERPs. Upper row: ERPs in the left frontal cluster of electrodes (highlighted in the topographical plot) depicting rehearsal effects from 0-500ms after the cue. Lower row: ERPs in the negative posterior cluster. (F) The ICS interaction effect as in Figure 2C calculated not based on single trials, but correlating condition-specific ERPs during item presentation with ERPs during cue presentation in each condition and every subject. This analysis revealed no significant results and an overall different pattern compared to the original analysis demonstrating that ERP effects cannot explain the reported ICS effects. (G) The exact same interaction analysis as in Figure 2C was repeated based on data with an alpha 8-20Hz band stop filter, to ensure that alpha power changes were not driving the effects. We obtained similar results as in the original analysis, contours highlight significant clusters (p_corr_–0.05).

**Figure S3:**
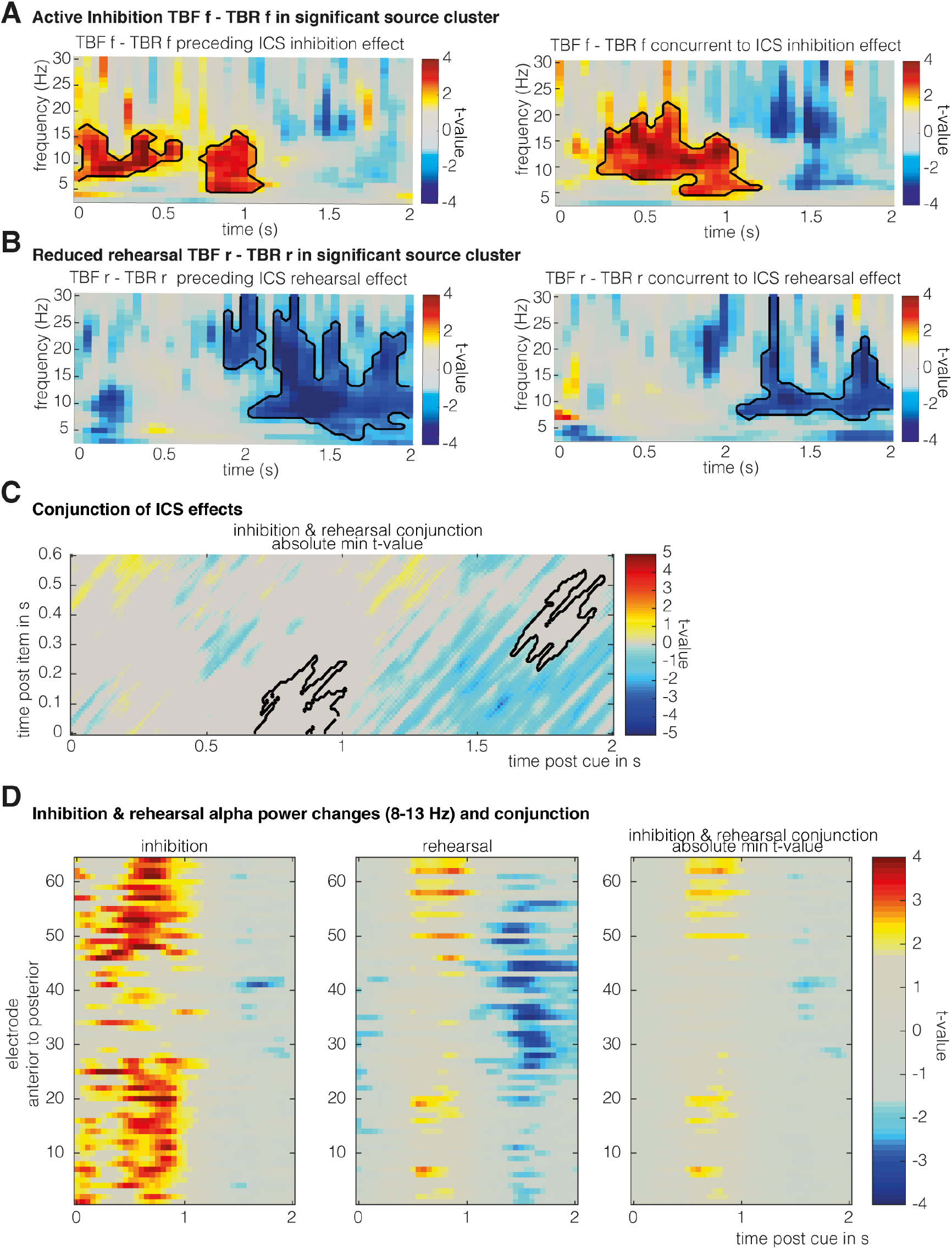
Time-frequency plots in significant source clusters and conjunction analyses related to Figure 3. (A) Time-frequency changes in the respective significant source clusters for the inhibition contrast (Figure 3 B&C). (B) Time-frequency changes in the respective significant source clusters for the rehearsal contrast (Figure 3 E&F), black contours highlight significant clusters (p<0.05). Please note that this analysis does not provide independent evidence since the clusters were defined by showing significant inhibition and rehearsal effects respectively. (C,D) Conjunction of inhibition and rehearsal effects. (C) Minimum t-statistic conjunction [S1,S2] of inhibition (TBF-f vs TBR-f) and rehearsal (TBF-r vs TBR-r) contrasts. For time points in which contrasts were in opposing directions, the conjunction was set to zero. An uncorrected threshold of p<0.05 was not exceeded at any time point. Contours show the positions of significant interaction clusters reported in Fig. 2C. (D) Alpha power changes (8-13Hz) in every electrode (sorted from posterior to anterior position). Plots depict t-values for the inhibition contrast (left), the rehearsal contrast (middle) and the minimum t-value conjunction (right).

**Figure S4:**
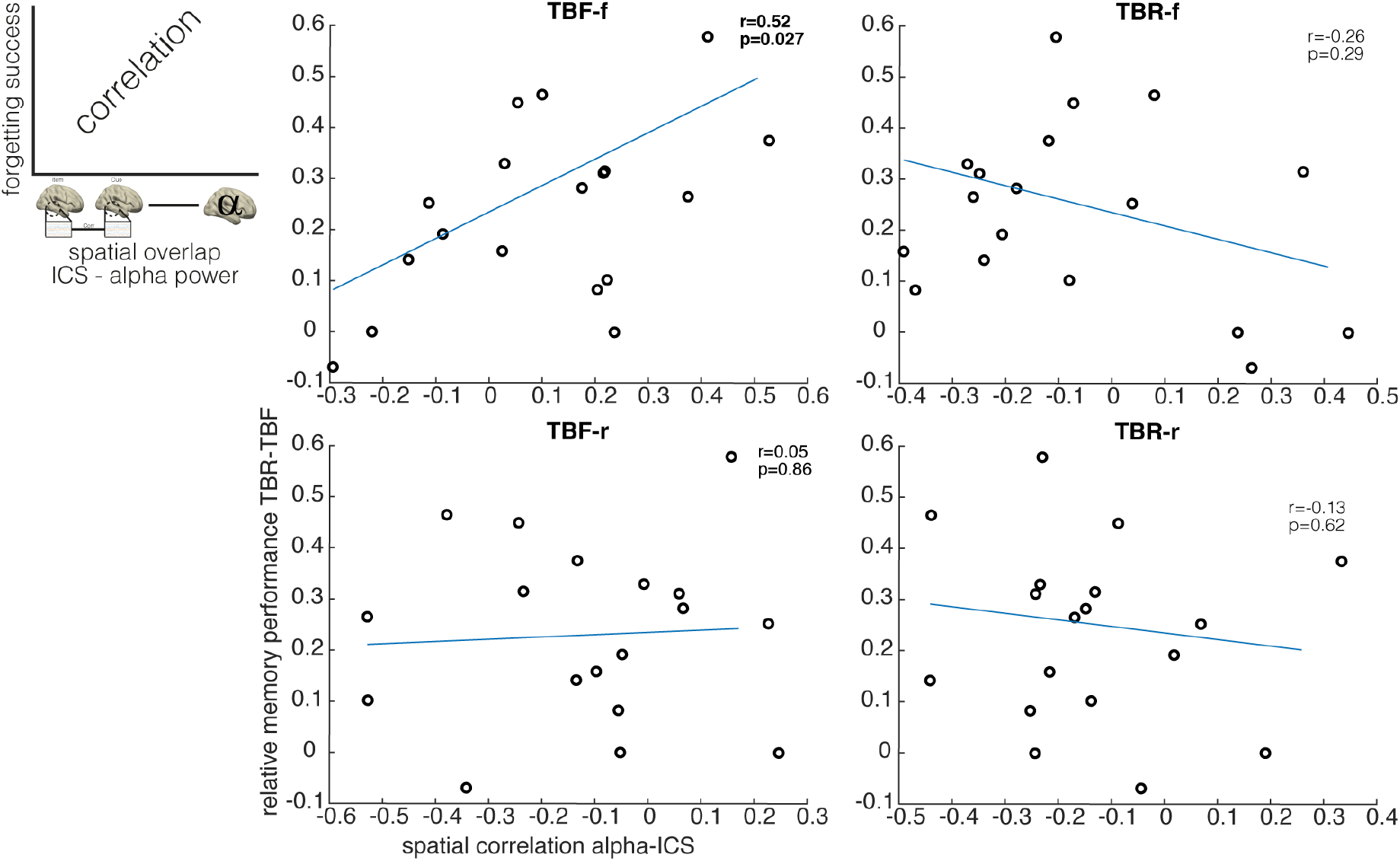
Overlap of alpha power changes and ICS predicts forgetting success related to Figure 4. To follow up on the finding that specifically alpha power increases preceding ICS inhibition effects are overlapping with ICS changes, we assessed whether the spatial overlap of early alpha power increases and ICS effects predicts forgetting success. The ICS-alpha overlap was evaluated by correlating mean alpha power source maps (0-500ms post cue, preceding ICS inhibition effects) to mean searchlight ICS maps (0-300ms item-time and 600-1000ms cue-time window), in every condition and subject. In order to investigate whether this alpha-ICS overlap predicts later forgetting, these alpha-ICS correlations were correlated with individual forgetting success (relative changes in TBR-TBF hit rate). A significant correlation was found only in the TBF-f condition, suggesting that voluntary forgetting depends on the spatial precision of early alpha power increases targeting item-specific representations.

## Notes

https://osf.io/upkwe/

### Competing Interest Statement

The authors have declared no competing interest.

### Summary of Updates

New analysis was added. Item-cue similarity analysis is now run on the source level using a searchlight approach. Supplemental figures are also updated incorporating additional control analyses.

